# The *Giardia* lamellipodium-like ventrolateral flange supports attachment and rapid cytokinesis

**DOI:** 10.1101/2021.01.31.429041

**Authors:** William R. Hardin, Germain C. M. Alas, Nikita Taparia, Elizabeth B. Thomas, Melissa Steele-Ogus, Kelli L. Hvorecny, Aaron R. Halpern, Pavla Tůmová, Justin M. Kollman, Joshua C. Vaughan, Nathan J. Sniadecki, Alexander R. Paredez

**Affiliations:** Department of Biology, University of Washington, Seattle, USA; Department of Mechanical Engineering, University of Washington, Seattle, USA; Department of Biochemistry, University of Washington, Seattle, USA; Department of Chemistry, University of Washington, Seattle, USA; Institute of Immunology and Microbiology, 1^st^ Faculty of Medicine, Charles University, Prague, Czech Republic; Department of Physiology and Biophysics, University of Washington, Seattle, USA; Bioengineering, University of Washington, Seattle, WA 98105, USA; Lab Medicine & Pathology, University of Washington, Seattle, WA 98105, USA; Center for Cardiovascular Biology, University of Washington, Seattle, WA 98105, USA; Institute for Stem Cell and Regenerative Medicine, University of Washington, Seattle, WA 98105, USA

## Abstract

Attachment to the intestinal epithelium is critical to the lifestyle of the ubiquitous parasite *Giardia lamblia*. The microtubule cytoskeleton plays a well characterized role in attachment via the ventral adhesive disc, whereas the role of the unconventional actin cytoskeleton is controversial. We identified a novel actin associated protein with putative WH2-like actin binding domains we named Flangin. Flangin complexes with *Giardia* actin and is enriched in the ventrolateral flange (VLF), a lamellipodium-like membrane protrusion at the interface between parasites and attached surfaces. Live imaging revealed that the VLF grows to ~1 μm in width after cytokinesis, then remains size-uniform in interphase, grows during mitosis, and is resorbed during cytokinesis. A Flangin truncation mutant stabilizes the VLF and blocks cytokinesis, indicating that the VLF is a membrane reservoir supporting rapid myosin-independent cytokinesis in *Giardia*. Rho family GTPases are important regulators of membrane protrusions, *Gl*Rac, the sole Rho family GTPase in *Giardia*, was localized to the VLF. Knockdown of Flangin, actin, and *Gl*Rac result in VLF formation defects indicating a conserved role for *Gl*Rac and *a*ctin in forming membrane protrusions, despite the absence of canonical actin binding proteins that link Rho GTPase signaling to lamellipodia formation. Flangin-depleted parasites challenged with fluid shear force in flow chambers had a reduced ability to remain attached, indicating a role for the VLF in attachment. This secondary attachment mechanism complements the microtubule based adhesive ventral disc, a feature that is particularly important during mitosis when the parental ventral disc begins disassembly in preparation for cytokinesis.

**Importance:** The ventrolateral flange (VLF) is a lamellipodium-like structure found at the host-parasite interface that has long been thought to be involved in parasite attachment. The proteins responsible for building the VLF have remained unidentified precluding manipulation of the VLF to determine its role in *Giardia* biology. We identified Flangin, a novel actin associated protein that localizes to the VLF, implicating *Giardia* actin in VLF formation. We demonstrate that: 1.) Flangin, actin, and *Gl*Rac are required for VLF formation, 2.) the VLF serves as a membrane reservoir to support *Giardia’s* incredibly fast cytokinesis, and 3) the VLF augments attachment, which is critical to parasitism. The microtubule-based adhesive ventral disc and the actin-based ventrolateral flange represent redundant means of maintaining attachment, the presence of redundant systems illustrate the importance of attachment to the lifestyle of this ubiquitous parasite.

## Introduction

The actin cytoskeleton is essential to maintaining cell shape and forms one of the central organizing scaffolds of all cells. Most of our understanding of actin regulation and function comes from studies performed in a limited number of model eukaryotes (1, 2). *Giardia lamblia* belongs to a potentially early diverging eukaryotic lineage, lacks all canonical actin-binding proteins (ABPs) used to regulate actin dynamics, including myosin, and has the most divergent eukaryotic actin identified to date (3–7). These unique attributes make *Giardia* an interesting model for studying novel mechanisms in regulating the assembly of actin microfilaments (8). Additionally, *Giardia* is a common waterborne pathogen that infects more than 200 million people each year, so its divergent actin cytoskeleton could be a potential therapeutic target (8, 9).

Despite the lack of canonical actin-binding proteins such as myosin, formin, cofilin, and the Arp2/3 complex, *Giardia* has a dynamic actin cytoskeleton that is required for conserved roles including membrane trafficking, cell shape, cell polarity, and cytokinesis (10, 11). Dynamic reorganization of *Giardia lamblia* actin (*Gl*Actin) during the cell cycle indicates the presence of actin regulatory proteins. A previous effort at biochemically isolating ABPs from *Giardia* identified several proteins that associate with *Gl*Actin. However, these proteins likely associate with monomeric actin, based the lack of filamentous localization patterns and the biochemical conditions used to isolate these proteins (12, 13). Therefore, the previously identified proteins are unlikely to account for the full complement of ABPs in *Giardia*.

Many ABPs have a small helical domain known as the Wasp Homology 2 (WH2) domain. This versatile domain is found in proteins that account for the most basic forms of actin regulation such as monomer sequestration, nucleation, capping, and depolymerization (14, 15). WH2 sequences are short 15-20 amino acid helices that bind to the hydrophobic cleft at the barbed (+) end of actin monomers, which avoids most cross strand interfaces of actin filaments permitting their use for diverse actin regulatory functions (16). *Giardia* actin is highly divergent with 58% average identity to conventional actins (yeast to man), yet the hydrophobic cleft is well conserved (Figure S1). Of the 147 residues that differ between mammalian skeletal actin and *Gl*Actin, only four variations fall within 5 Å of the WH2 binding site. Two are conservative hydrophobic substitutions (Ile to Val, residues 341 and 345). The side chain of the third, Ala to Ser at position 144, does not interact with the WH2 domain. The fourth substitution, Ser to Ala at position 350, causes Gln437 on the WH2 domain to lose a hydrogen bonding partner. However, this residue is not considered part of the core residues conserved within WH2 domains (16), suggesting that WH2 or WH2-like domains could regulate actin in *Giardia*.

Here we used a bioinformatics approach to identify putative WH2-like domain containing proteins and identified Flangin, a protein that localizes to *Giardia’s* lamellipodium-like ventrolateral flange (VLF). The VLF is a thin flexible membrane protrusion encircling the ventro-lateral periphery of trophozoites (life stage that colonizes the intestine). Although morphologically similar to lamellipodia, a major difference is that the VLF is not known to have any role in motility (17–19). Based upon appearance, it has been suggested that the VLF plays a role in parasitic attachment (20, 21). The only experimental support for the VLF participating in attachment comes from a study where the ventral disc was mechanically impeded from establishing suction pressure with a microfabricated surface; a small subset of cells still managed to attach with “adhesive” contacts between the VLF and the microfabricated surface (22). Molecular components of the VLF were unknown preventing further testing of the VLFs role in attachment.

Our identification of a VLF component presented the opportunity to explore the role of the VLF in attachment as well as determine how a lamellipodia-like membrane protrusion is assembled in the absence of canonical ABPs such as the Arp2/3 complex. We use a combination of live and fixed cell imaging to investigate VLF and Flangin dynamics. We demonstrate that Flangin, *Gl*Actin, and *Giardia’s* sole Rho family GTPase, *Gl*Rac, all have roles in VLF formation. With the ability to perturb VLF formation for the first time, we perform attachment assays under fluid flow and confirm that the VLF has a role in mediating parasite attachment. This finding is important because most studies aimed at understanding *Giardia* attachment have focused on the suction-cup like microtubule based adhesive ventral disc (7, 23). Our findings indicate that the ventral disc is only part of the attachment mechanism. The importance of *Gl*Actin in VLF formation and overall cellular organization highlights the importance of the actin cytoskeleton in *Giardia* attachment, which was previously controversial (7).

## Results

### *Giardia* possesses a lamellipodium-like structure that contains *Gl*Actin and Flangin

To identify putative actin-binding protein(s), we performed a bioinformatics search of the *Giardia* genome for WH2 domain containing proteins. Using PHI blast and various PHI patterns (see materials and methods), 216 proteins were identified that contain one or more putative WH2-like domains (Table S1). Our search parameters were low stringency; while we likely have a high false positive rate, the comprehensive list is a valuable starting point for this and future studies. Since *Giardia* lacks Arp2/3 complex and formin actin nucleators, we were particularly interested in proteins with multiple WH2-like domains. Such proteins could potentially have a role in actin nucleation, which is critical to cytoskeletal regulation (16, 24). Our search yielded eight proteins with multiple predicted WH2 domains. One protein, GL50803_7031, hereafter referred to as Flangin was particularly intriguing because in addition to three putative WH2-like domains, this protein also possesses a Bro1 domain (Interpro E=2^-13^) at the N-terminus that is implicated in F-actin binding (Figure S2). The mammalian protein Alix functions as a regulator of membrane remodeling and it possesses a Bro1 domain that has been shown to directly bind F-actin (25, 26). Whether F-actin binding is a general feature of Bro1 domains has yet to be determined. Due to the presence of a putative F-actin binding domain and three WH2-like domains that potentially modulate actin function, we proceeded to test for interaction with *Gl*Actin. Immunoprecipitation of triple hemagglutinin epitope tagged Flangin (Flangin-3HA), pulled down *Gl*Actin versus a wild type negative control indicating molecular association of Flangin and *Gl*Actin (Figure 1A).

**Figure 1.**
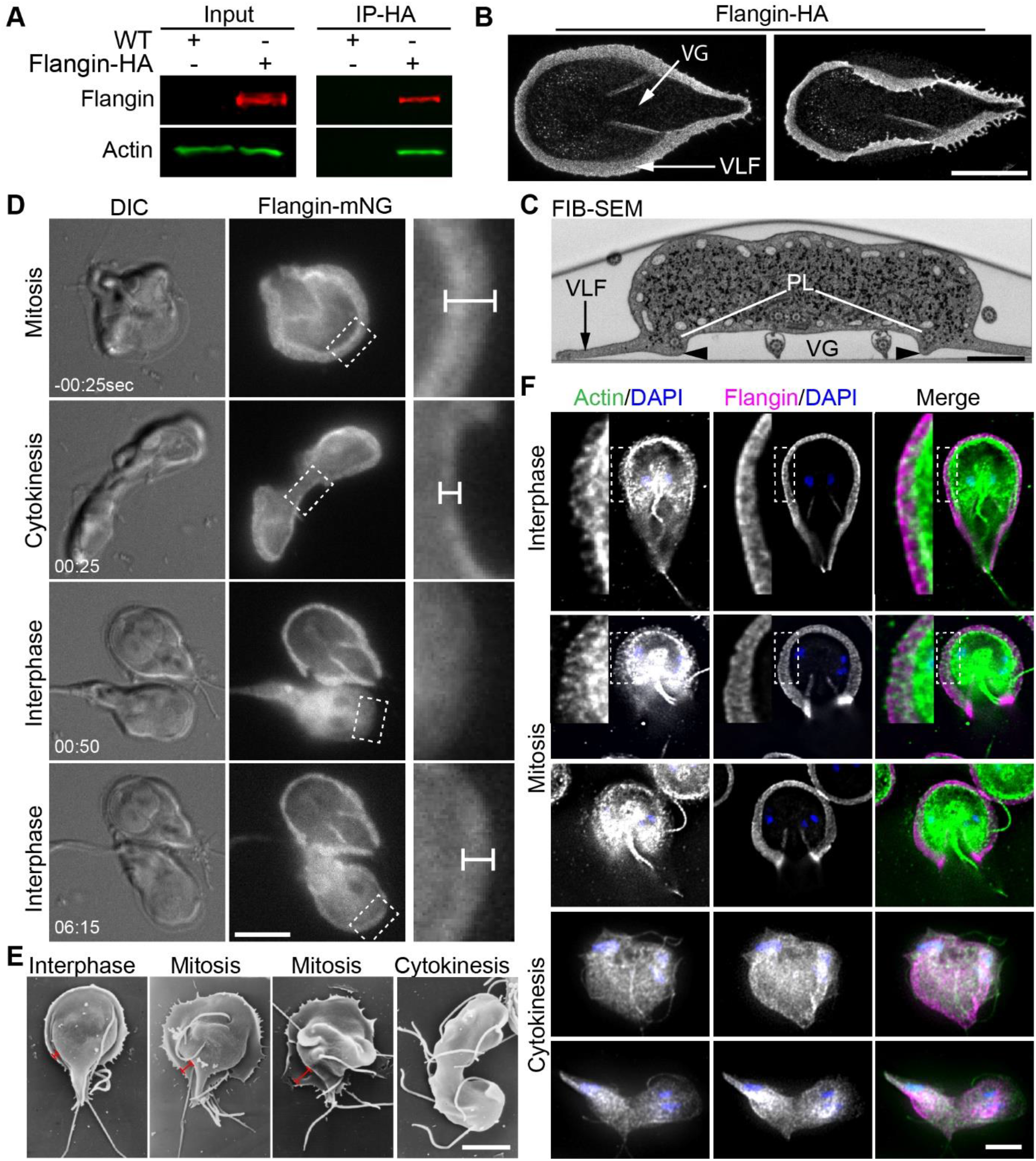
*Giardia* possesses a lamellipodium-like structure that contains *Gl*Actin and Flangin. (A) Immunoprecipitation of Flangin-HA from cell extracts, followed by anti-*Gl*Actin Western blotting demonstrates Flangin associates with *Gl*Actin. (B) Maximum projections of expanded trophozoites imaged with confocal fluorescence microscopy. Flangin-HA marks the VLF and membrane protrusions associated with the cytoplasmic portions of the posterolateral flagella axonemes that bound the ventral grove (VG). The left image shows that the VLF can completely encircle the base of the cell and the image on the right shows that the VLF is thin and flexible. (C) Transverse section of wild-type *Giardia* (posterior to the ventral flagella exit point) as viewed with Focused Ion Beam-Scanning Electron Microscopy (FIB-SEM). Arrowhead marks the membrane protrusion associated with Flangin and the cytoplasmic axonemes of the posterolateral (PL) axonemes that bound the ventral groove. (D) Live cell imaging of Flangin-mNG during cytokinesis. Compared with interphase cells the VLF is wider in mitosis and was resorbed during cytokinesis. Upon cytokinesis, the VLF disassembled and Flangin-mNG translocated to the cytoplasm. Daughter cells lack a VLF, but assembly was initiated almost immediately after division. Flangin-mNG was recruited back to the VLF as it reformed(see Movie 1 and 2). Magnified views of the boxed area show Flangin is at the leading edge of the VLF. Dimension bars in C and D point out VLF width. (E) Scanning electron microscopy shows VLF growth during mitosis and resorption during cytokinesis. (F) Immunofluorescence localization of *Gl*Actin (green), Flangin-HA (magenta), and DNA (blue), throughout the cell cycle (see Figure S3 for tubulin staining). *Gl*Actin and Flangin accumulate in the VLF during interphase and mitosis. The insets show a magnified view of *Gl*Actin and Flangin in the VLF. During cytokinesis VLF localized Flangin and *Gl*Actin translocated to cytoplasm as the VLF is disassembled. Note that the interphase and mitotic cells are partial projections optimized to show *Gl*Actin localization, while the entire Z-stack was projected for the cells in cytokinesis to show that the VLF has been resorbed. Scale bars= 5 μm except (C)=1 μm.

Flangin-3HA was then localized with expansion microscopy (27, 28), a super-resolution microscopy method which provided ~70 nm spatial resolution here, or roughly four times the resolution of conventional confocal microscopy. In *Giardia* trophozoites, Flangin-3HA was found to localize to the ventrolateral flange, a thin flexible membrane protrusion surrounding the whole cell with a gasket-like appearance (VLF; Figure 1B). In addition to localization at the VLF, Flangin was found along the boundaries of ventral groove, which is demarcated by the cytoplasmic portions of the posterolateral flagella axonemes (Figure 1B, C).

We recently developed methods to film fluorescently tagged proteins through the *Giardia* cell cycle (11). To follow Flangin dynamics over the cell cycle, we tagged Flangin with mNeonGreen (Flangin-mNG) and performed live cell imaging. Relative to interphase, the VLF width increased by 69% in mitosis (0.93 ± 0.23 μm to 1.57 ± 0.44 μm n=18). VLF expansion begins in anaphase so that the expanded VLF is in place when the ventral disc begins disassembly at the anaphase/telophase transition (Figure S3A (11, 29, 30)). Cytokinesis is initiated after nascent ventral disc assembly (10, 29, 30). At the onset of cytokinesis, Flangin re-localized from VLF to the cytoplasm as the VLF was resorbed (Figure 1D, Movie 1). Note that *Giardia* cytokinesis involves detachment in order for daughter cells to swim in opposite directions, which generates the membrane tension required for furrow progression and abscission (11, 30, 31). Within 60 seconds of completing cytokinesis, daughter cells initiated VLF protrusion at the anterior of the cell (Movie 2). It should be noted that the cell body bulges above the VLF (see Figure 1C), which slightly impinges on our ability to see and measure initial VLF growth with differential interference contrast (DIC) microscopy used for the measurements in this paper. After VLF assembly is initiated at the anterior, VLF protrusion propagated along the cell with the posterior VLF being the last to complete growth. Daughter cells noticeably elongate and restore their typical shape during posterior VLF expansion (Movie 2). On average, the anterior portion of the VLF reached a maximum width of 0.97±0.079 μm after 155±32 seconds with a protrusion rate of 0.39±0.1 μm/minute (n=20), which is near the protrusion rate of lamellipodia (32, 33). Unlike lamellipodia, which continuously grow and shrink to drive cell motility, the VLF grew without apparent shrinking or retracting before reaching the final interphase width (Movie 1, 2). Consistent with previous observations (30), scanning electron microscopy of wild-type trophozoites verified that the VLF grows in width during mitosis and is resorbed during cytokinesis (Figure 1E).

Given the morphological similarity of the VLF to lamellipodia and the biochemical association of Flangin with *Gl*Actin, we assessed *Gl*Actin presence in the VLF. We imaged actin in fixed cells using a *Gl*Actin specific antibody because it is not possible to fluorescently tag *Gl*Actin or use conventional live actin reporters such as LifeAct or SiR-Actin (10). Both filamentous *Gl*Actin and Flangin-3HA, were found throughout the VLF in interphase and mitotic trophozoites (Figure 1F). Additionally, Flangin was observed in association with the ends of *Gl*Actin filaments in the cell body (Figure S3B). Together, our results indicate that Flangin and *Gl*Actin are biochemically associated and are components of the VLF.

### Flangin and *Gl*Actin are necessary for VLF assembly

To reveal if Flangin and *Gl*Actin are required for VLF formation, we depleted these proteins and measured the impact on the width of the VLF. Translation-blocking antisense morpholinos were used to deplete *Gl*Actin, Flangin, or human beta-globin as a negative control. Quantitative western blotting revealed ~60% depletion of *Gl*Actin at the population level, 24 hours after morpholino treatment (Figure S4A). Knocking down actin, in addition to previously reported cellular organization defects (10), resulted in either an inverted or narrow VLF phenotype compared to the negative morpholino control (Figure 2A). For cells with the inverted phenotype, the VLF was not visible by DIC microscopy and Flangin was found to be localized around the edge of the cell body (23.2% of cells, n=73). In the remainder of cells with the narrow VLF phenotype, the width of the anterior VLF was significantly decreased (Figure 2A and 2B). An antisense-translation blocking morpholino targeting Flangin achieved ~60% knockdown (Figure S4B). Flangin-depleted cells had a narrow VLF phenotype when compared to the control morpholino treated cells (Figure 2A and 2B). However, during mitosis when the VLF grows in width, we were able to find Flangin-depleted cells with expanded VLFs where *Gl*Actin but not Flangin was found at the leading edge of the VLF (Figure 2C). These results are consistent with a role for *Gl*Actin in driving membrane protrusion and a role for Flangin in stabilizing the VLF.

**Figure 2.**
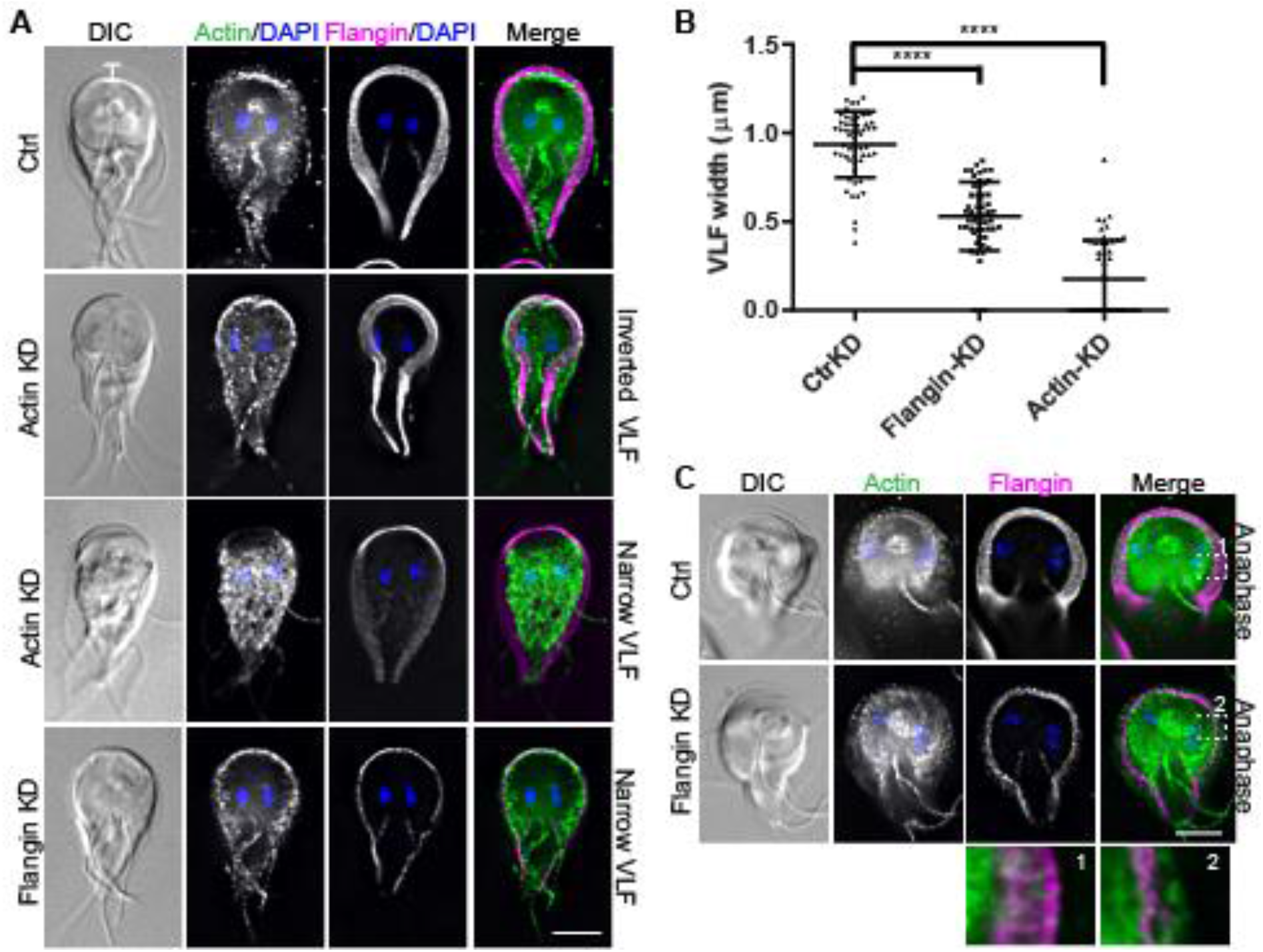
Flangin and *Gl*Actin are necessary for VLF assembly. (A) Trophozoites were stained for *Gl*Actin (green), Flangin-HA (magenta), and DNA (blue). *Gl*Actin knockdown (KD) resulted in inverted and collapsed VLF morphology. Flangin-HA KD similarly resulted in a thin VLF phenotype. (B) Quantification of VLF width when measured at the cell anterior from three independent experiments; control (n=57), Flangin-HA (n=54), and actin (n=51). Statistical significance was evaluated for Flangin-KD and Actin-KD respectively, t test. ****, P<0.0001. (C) Mitotic Flangin depleted cells were capable of extending their VLF beyond the leading edge of Flangin. The insets show actin beyond the leading edge of Flangin-HA in Flangin-HA KD cells during mitotic VLF extension. Scale bars= 5 μm.

### Flangin is a stable component of the interphase VLF

In contrast to lamellipodia, which are continuously remodeled to drive cell motility, VLF size under our imaging conditions is static after initial re-growth in interphase (Movie 2). We postulate that the VLF could maintain a fixed width either through a balance of disassembly and assembly processes or the use of a static scaffold. Without a live *Gl*Actin marker, it is not currently possible to assess actin dynamics. To probe whether Flangin is a dynamic or stable component of the assembled VLF, we assessed Flangin dynamics with fluorescence recovery after photobleaching (FRAP). FRAP of Flangin-mNG revealed minimal recovery over a 12-minute window (n=35 cells). We conclude that Flangin is a stable structural component of the VLF (Figure 3A and 3B).

**Figure 3.**
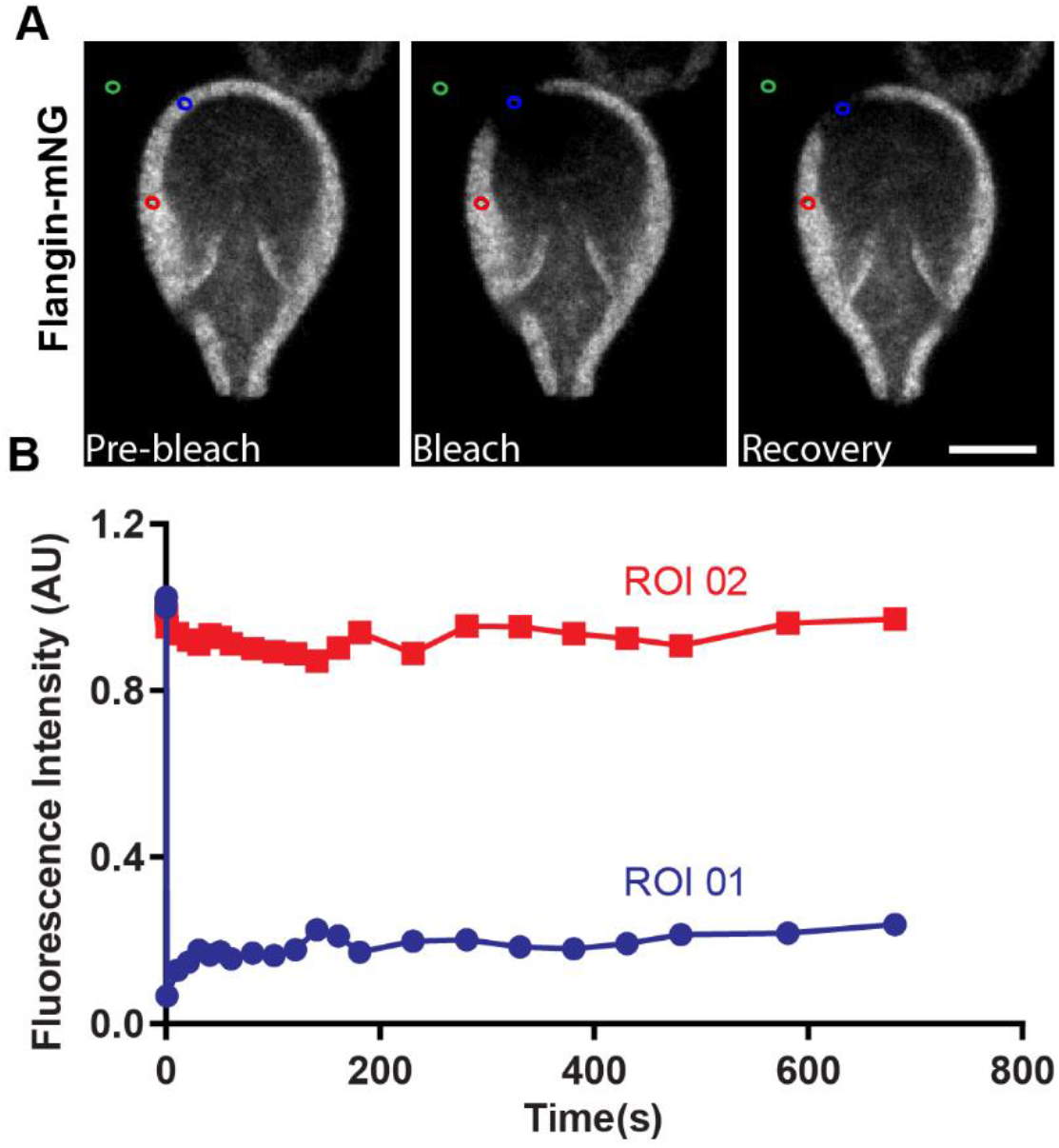
Flangin is part of a stable structure. (A) Fluorescence Recovery after Photobleaching (FRAP) was performed on the VLF of Flangin-mNG cells, Blue circle bleached ROI 01, Red circle non-bleached ROI 02, and Green circle background ROI 03. (B) ROI 01 showed minimal post-bleaching recovery after 12min. Similar results were observed for 35 other FRAP experiments including the small membrane protrusion associated with the posterolateral axonemes. Scale bar= 5 μm.

We also tested whether the N-terminal Bro1 domain (amino acids 1-362) or C-terminal WH2 region (amino acids 301-995) of Flangin are sufficient to localize the protein to the VLF. The N-terminal Bro1 domain is sufficient to recruit Flangin to the VLF whereas the C-terminal fragment was distributed through the cell body without enrichment at the VLF (Figure S5). It is important to note that Bro1 domains are boomerang shaped membrane binding proteins that could have a role in inducing membrane curvature or sensing curvature similar to BAR domains, which are not found in *Giardia* (34–36). The Bro1 domain is sufficient for recruitment to the VLF, but whether this is mediated by interaction with specific lipids, interacting proteins, or the unique membrane curvature of the VLF remain to be determined.

### VLF membrane is a reservoir to support rapid cytokinesis

*Giardia* trophozoites are amongst the fastest dividing eukaryotes, requiring a median time of just 50 seconds to complete cytokinesis (11). This timing does not include breaking of the fragile cytoplasmic bridge that sometimes persists from seconds to tens of minutes (30). Division has a concomitant need for increased plasma membrane; for a typical eukaryotic cell, this involves an approximately 1.5 fold increase in surface area (37). We previously proposed that *Giardia* might possess a reservoir of membrane to help support this rapid division (11). In model eukaryotes, membrane protrusions have been shown to act as surface membrane reservoirs to support cell division (38, 39). The observation that the VLF is resorbed during cytokinesis suggests that the VLF could contribute membrane for cytokinesis. To test for a role in cytokinesis, we filmed Flangin-depleted cells using four-dimensional (4D) DIC microscopy (Figure 4A, Movie 3, 4).

**Figure 4.**
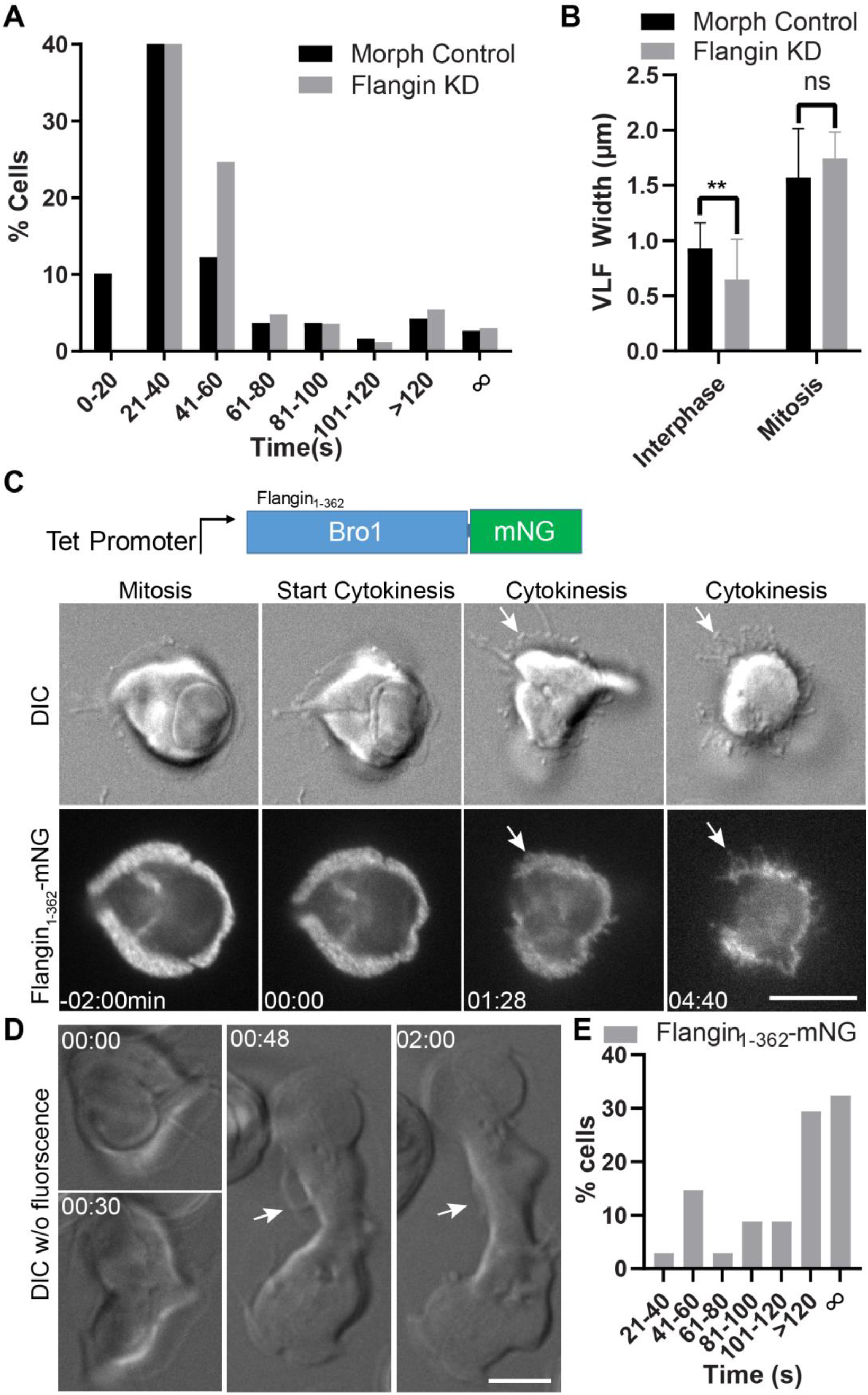
VLF breakdown is necessary for cleavage furrow progression. (A) Histogram showing the timing of cytokinesis for morpholino control (black) and Flangin KD cells (gray). See Movies 3 and 4). Cytokinesis median time: control 31 sec (n=188) and Flangin KD 35 sec (n=166). Due to sampling frequency (timing between image stacks) the difference in median values in this experiment is not meaningful, similar to our previous study, more than 90% of the cells in both groups complete cytokinesis by 120 seconds indicating that there is no cytokinesis defect. (B) Flangin KD cells grow their VLF before cytokinesis similar to control cells (See Movie 4). (C) A tet inducible N-terminal truncation of Flangin interferes with VLF retraction during cytokinesis. This blocks furrow progression and the ability of the parasite to lift off the cover glass (Compare Movie 1 and 5). White arrow points to regions of the VLF that should have been resorbed by this point in cytokinesis. (D) DIC imaging of the same cell line without fluorescence. This is an example of a moderate defect where the VLF persists and cytokinesis took 8 minutes to complete (see Movie 6). (E) Cytokinesis timing for Flangin1-362-mNG filmed with DIC only (n=34). Scale bar= 5 μm.

We did not observe delays or defects in cytokinesis when Flangin was depleted. Our previous analysis of *Giardia* cytokinesis found that 89% of wild type cells complete cytokinesis within two minutes with a median time of 50 seconds (11). Here, more than 90% of morpholino control or Flangin-depleted cells completed cytokinesis in under two minutes. The median times for cytokinesis was 31 seconds for the morpholino control (n=188) and 35 seconds for the Flangin-depleted cells (n=166). Although the four second difference is statistically significant, in the context of our previous results and slight differences in image acquisition parameters for individual time lapse movies, the small difference in timing represents a sampling issue (number of optical sections in each image stack and interval between image stacks changes sampling) and is not biologically meaningful.

The observation that Flangin-depleted cells can restore their VLFs during mitosis accounts for the lack of cytokinesis defects. Consistent with Figure 2C, Flangin-depleted cells with narrow VLF widths were able to expand their VLFs in mitosis (Movie 4). We found that Flangin-depleted cells increased their VLF width during mitosis by about three-fold (Figure 4B). There was no significant difference in the maximum VLF width before cytokinesis for Flangin-depleted cells (1.76±0.25 μm) and morpholino control cells (1.57 ± 0.44). Thus, Flangin-depleted cells effectively establish a membrane reservoir before initiating cytokinesis.

While Flangin depletion did not impair cytokinesis, the tetracycline inducible Flangin_1-362_-mNG fusion protein had dominant negative function that impaired cytokinesis (Figure S5). Live imaging of Flangin_1-362_-mNG revealed that the truncation protein impairs cytokinesis by stabilizing the VLF. We observed two types of cytokinesis defects: complete failure or partial furrow progression (n=21) (Figure 4C, Movie 5). The example in Figure 4C is the more severe phenotype of complete failure to divide where stabilized regions of the VLF result in filipodia like projections seen in six cells. The other 15 cells had some furrow progression, where portions of the VLF did not disassemble, resulting in failed cytokinesis where one emerging daughter was smaller due to an effectively reduced membrane reservoir.

When filming fluorescently tagged proteins, we sometimes observed mitotic cells that stall in cytokinesis or never initiate the process after mitosis, likely a result of oxidative stress (40). Therefore, we examined this cell line using long-term DIC imaging to avoid oxidative stress. Under DIC imaging, the cytokinesis failure rate of wild type and control cells is between 2-3% (11, 41). When Flangin_1-362_-mNG was filmed with 4D DIC imaging, 52.9% of cells took more than 2 minutes to divide and 32.3% failed to complete cytokinesis (n=34, Figure 4D, 4E, Movie 6), confirming that the N-terminal fragment interferes with the dynamics of the VLF and cytokinesis. Taken together, our imaging results illustrate a requirement for disassembly of the VLF for cytokinesis.

### *Gl*Rac has a conserved role in mediating actin-based membrane protrusions

Rho family GTPases are important regulators of actin-based membrane protrusions in model eukaryotes (42). We questioned whether *Gl*Rac (GL50803_8496), *Giardia’s* sole Rho family GTPase, might also regulate VLF formation. N-terminally HA-Tagged *Gl*Rac localized to the VLF with some enrichment near *Gl*Actin filaments (Figure 5A). Live imaging of Halo-*Gl*Rac, which avoids the effect of detergents on Rho GTPase localization (43), indicated a more uniform distribution of *Gl*Rac in the VLF. Since Rho GTPase activity is regulated by nucleotide state, we sought to determine if *Gl*Rac is actively signaling at the VLF. CRIB/PBD domains are specifically recruited to active GTP-loaded Rho GTPases and have been used as the basis for Rho GTPase signaling biosensors (44, 45). PAK kinase (GL50803_2796) contains the only CRIB/PBD domain in the *Giardia* genome. We tagged the CRIB/PBD domain of *Giardia’s* PAK kinase with mNG to create a CRIB-mNG based *Gl*Rac signaling biosensor. CRIB-mNG was robustly recruited to the VLF, suggesting a role for active *Gl*Rac in regulating VLF assembly (Figure 5B).

**Figure 5.**
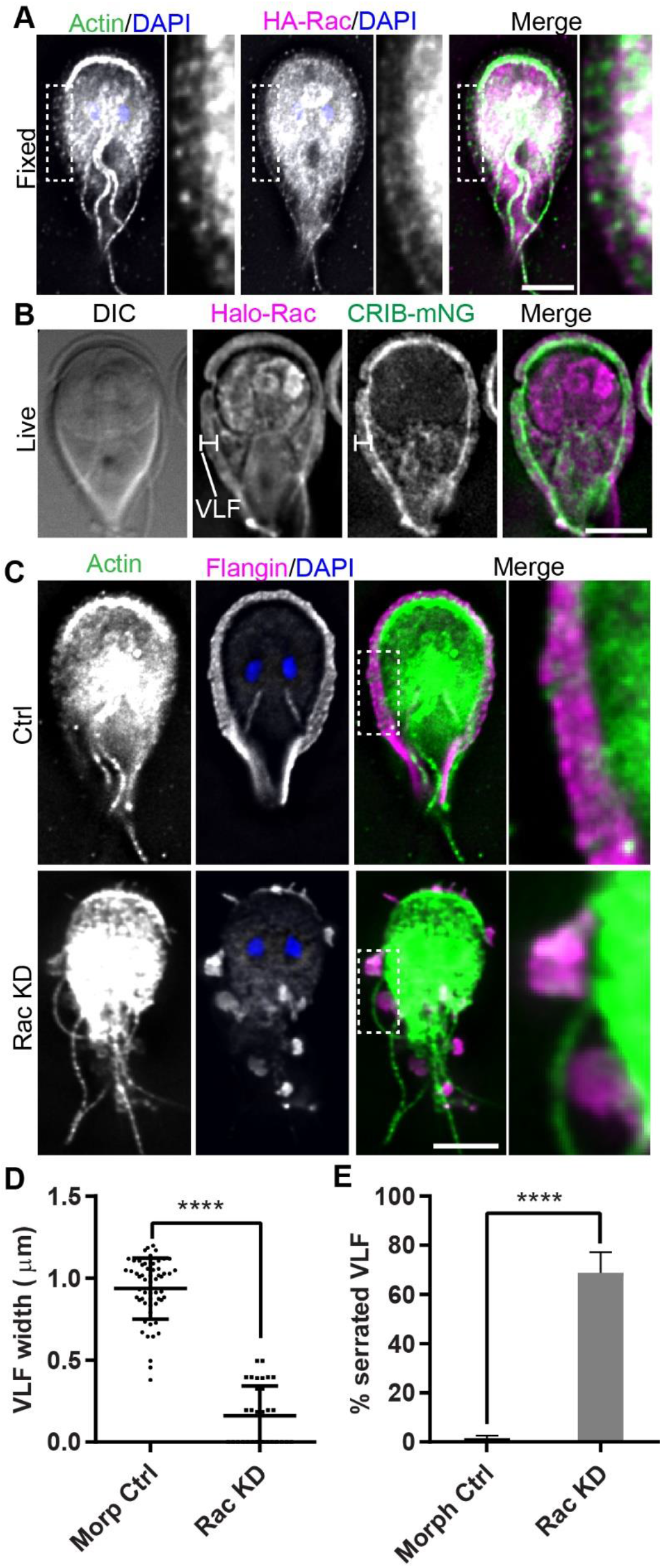
*Gl*Rac signaling is required for VLF assembly. (A) Immunofluorescence localization of *Gl*Actin (green), HA-Rac (magenta), and DNA (blue). *Gl*Actin and *Gl*Rac localize to the VLF. (B) Live cell imaging of Halo-Rac (magenta) and CRIB-mNG (green) indicate that *Gl*Rac is actively signaling in the VLF. (C) *Gl*Rac KD resulted in serrated VLF’s not observed in control cells. *Gl*Actin and Flangin localize to the remaining sporadic VLF protrusions. (D) Quantification of VLF width measured at cell anterior from three independent experiments: control (n=57) and *Gl*Rac KD (n=32). Statistical significance was evaluated using the t-test. ****, P<0.0001. (E) Quantification of serrated cells. Cells with three or more VLF gaps were defined as serrated; control (n=596) and *Gl*Rac KD (n=618). Statistical significance was evaluated using the t-test. ****, P< 0.0001. Scale bars=5 μm.

To determine if *Gl*Rac signaling is necessary for VLF formation, we depleted *Gl*Rac with translation blocking morpholinos. *Gl*Rac was depleted by ~70% compared to the morpholino control 24 hours after morpholino treatment (Figure S4C). *Gl*Actin and Flangin-HA localization was extremely perturbed in *Gl*Rac knockdown cells. *Gl*Actin and Flangin were found to accumulate in patchy VLF protrusions resulting in a serrated appearance (Figure 5C). Quantification of the *Gl*Rac knockdown phenotypes indicated a significant decrease in the anterior VLF width and a significant number of cells with a serrated phenotype that was not observed in control cells (Figure 5D and 5E). We interpret these results to indicate that Flangin and *Gl*Actin recruitment is downstream of *Gl*Rac signaling and that the role of Rho GTPases regulating membrane protrusions is broadly conserved.

### The VLF contributes to attachment

We depleted Flangin, *Gl*Actin, and *Gl*Rac to test their role in attachment. Initially, a basic attachment assay was performed where the proportion of trophozoites that could adhere to the side of culture tubes was determined. In this assay, knockdown of Flangin did not disrupt attachment. Earlier studies of *Giardia* attachment found that cells could attach under conditions where the ventral disk could not be employed (22). Likewise, disruption of ventral disc organization is not sufficient to disrupt attachment (46, 47). Defects are only observed after challenge assays, which is consistent with redundant means of attachment. We did observe fewer *Gl*Rac- and *Gl*Actin-knockdown cells attached to the culture tube compared to the control (Figure 6A). The reduced attachment of *Gl*Rac- and *Gl*Actin-depleted cells likely result from gross disorganization of the cytoskeleton (10) that results in combined disruption of the VLF and ventral disc.

**Figure 6.**
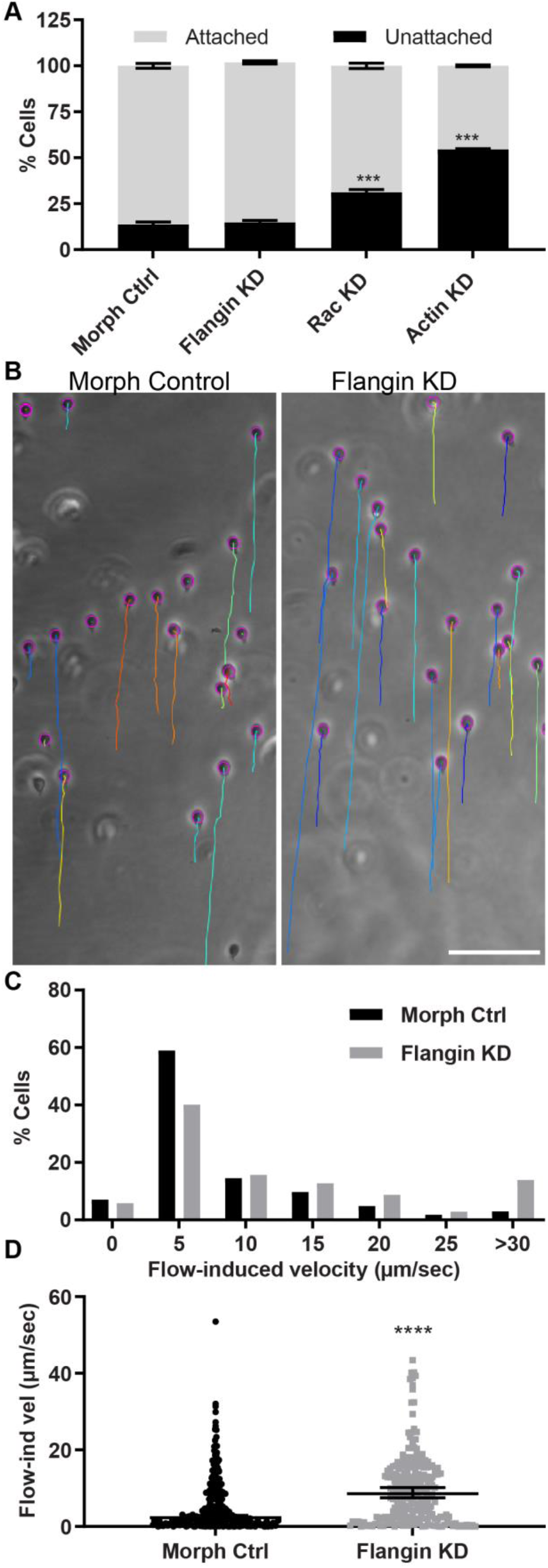
The VLF has a role in attachment. (A) Quantifying the role of Flangin, *GlRac*, and actin in non-challenged cell attachment. Each protein was depleted and then attached and unattached cells were counted. Three technical replicates for unattached cells were conducted. Mean ± SEM n=3, control 13.73 ± 0.78, Flangin 14. 81 ± 0.62, *Gl*Rac 31.23 ± 0.83, and *Gl*Actin 54.48 ± 0.25. *Gl*Rac and *Gl*Actin had statistically significant reduction in their ability to attach versus the morpholino control, t-test ***, P< 0.001. (B) Flow chamber assay to test the role of the VLF in attachment. A flow rate of 100ul/s induced sliding. The paths of cells were followed with TrackMate, the paths at the end of the 10s challenge are indicated by the colored lines. See Movie 7. (C) Histogram of flow induced velocities for the standard morpholino control n=268 and anti-Flangin morpholino treated cells n=248 analyzed from six challenge assays. (D) Plot of measured values error bars are median with 95% confidence interval; statistical significance determined with a Mann-Whitney U test. ****, P<0.0001. Scale bar= 100 μm.

Since Flangin depletion disrupts VLF maintenance but not overall cytoskeletal organization, Flangin knockdown permits direct assessment of the VLF’s role in attachment. We used flow chambers to challenge morpholino control and Flangin-knockdown cells with fluid shear force (Figure 6B, Movie S4). In this assay, Flangin-depleted cells were more frequently displaced and moved at higher flow induced velocities than control morpholino treated cells (Figure 6C, 6D). This result is consistent with the VLF being an auxiliary mediator of attachment.

## Discussion

### Flangin and *Gl*Actin are the first structural components of the VLF to be identified

We identified Flangin, a protein with three putative WH2-like domains and a Bro1 domain all implicated in mediating interaction with actin. While *in vitro* assays testing direct interactions with actin are routine in model eukaryotes, it has proven challenging to produce sufficient quantities of *Gl*Actin for these assays (10, 12). Therefore, we cannot yet conclude whether the WH2-like and/or Bro1 domains of Flangin directly mediate actin interaction. Yet, immunoprecipitation of HA-Flangin indicates association with *Gl*Actin. Flangin and *Gl*Actin were localized to the VLF, establishing these proteins as the first identified components of the VLF.

The reduction in VLF width observed in *Gl*Actin- and Flangin-depleted cells indicates that these proteins are integral to the formation and maintenance of the VLF. Live imaging revealed that Flangin is recruited to the VLF concomitantly with VLF formation after cell division (Figure 1C, Movie 1). FRAP studies indicate that once in place, Flangin is part of a stable structure (Figure 3), which is consistent with the lack of dynamic growing and shrinking that occurs in lamellipodia. Cells with a collapsed VLF likely reflect a stronger phenotype and more clearly highlight the critical role actin plays in driving VLF protrusion (Figure 3A and B). Live imaging revealed that the VLF protrudes from newly divided cells at a protrusion rate of ~0.4um/min until it reaches ~1 μm in width. Although Giardia is missing proteins involved in cell motility, the protrusion rate is on the same order of magnitude as cells using lamellipodia dependent mesenchymal motility (32, 33), suggesting that the *Gl*Actin cytoskeleton remains fully functional. Indeed recent observations unexpectedly demonstrated that *Giardia* may be capable of phagocytosis, which is an actin-dependent process that requires the formation of large membrane protrusions (48).

Like *Gl*Actin depletion, Flangin-depletion resulted in reduced VLF width, but during mitosis the VLF can extend beyond the position of Flangin (Figure 2C). We propose that *Gl*Actin is necessary for VLF membrane expansion, whereas Flangin is required for persistence of the protrusion. Tropomyosin, a coiled-coil protein that winds around and stabilizes actin filaments, has been shown to enhance the persistence of lamellipodial protrusions in fibroblasts (18). By analogy, Flangin may have a role in stabilizing *Gl*Actin by either binding along its length or capping filaments. Co-localization of *Gl*Actin and Flangin suggests that Flangin is associated with actin filaments (Figure 1E, 2C, and 5C). Re-sliced image stacks revealed that Flangin associates with the ends of filaments (Figure S3B). The development of a live *Gl*Actin marker or *in vitro* polymerization assays will be necessary to formally test how Flangin impacts *Gl*Actin dynamics.

### *Gl*Rac signaling is critical for VLF assembly

Rho family GTPase regulate lamellipodia formation through coordination of the actin cytoskeleton (49, 50). We found *Gl*Rac and its corresponding signaling biosensor CRIB-mNG were both recruited to the VLF (Figure 5A, 5B). *Gl*Rac depletion resulted in a serrated VLF phenotype (Figure 5C, 5D, and 5E). These findings indicate that *Gl*Rac is an essential upstream regulator of VLF formation and parallels the role of Rho family GTPases regulating actin and membrane protrusions in eukaryotes with conventional ABPs (51). That *Gl*Actin and *Gl*Rac are necessary for VLF formation raises the interesting possibility that the VLF has a shared origin with lamellipodia. Homologous organelles with a common origin can evolve new functions. Whether the VLF’s resemblance to lamellipodia stems from a common origin or convergent evolution remains an intriguing question.

### Two Critical Roles for the VLF in *Giardia*

In contrast to dynamic lamellipodia that drive cell motility, the VLF is only dynamic during initial growth and then again during mitosis and cytokinesis. Our functional assays indicate that the VLF is required for attachment of trophozoites (Figure 6 B-D). The increase in VLF width during mitosis is a remarkable innovation. After the initiation of cytokinesis, the ventral adhesive disc is disassembled to re-direct components to the nascent daughter ventral discs and also to clear a path for furrow progression (Figure S3) (11, 30). Trophozoites remain attached until the initiation of cytokinesis, after which the trophozoites detach and swim apart to generate membrane tension for furrow progression and abscission (11, 30). To our knowledge, no eukaryote can complete cytokinesis as quickly as *Giardia*. The incredibly fast cytokinesis is likely the result of evolutionary pressure to minimize the amount of time cells spend detached, which avoids being swept down the intestinal tract. We propose that the increased size of the VLF during mitosis serves to maintain attachment when the parental ventral disc is being disassembled and expected to lack full function. On smooth surfaces the VLF could contribute to attachment by maintaining suction pressure under the cell, but the VLF could also have adhesive properties as suggested by Erlandsen and others (22). The presence of redundant attachment mechanisms highlights the importance of intestinal attachment to the *Giardia* lifecycle. Our study also brings to light that *Giardia* attachment studies should consider contributions of both the ventral disc and the VLF.

We also propose that the VLF is a surface membrane reservoir to support rapid cytokinesis. The use of a reservoir to support specific events during cell-division has already been observed in *Giardia*. Specifically, the median body serves as a tubulin reservoir for both spindle and daughter ventral disc assembly (11, 46, 52). The growth of the VLF during mitosis would provide a larger pool of surface membrane to support rapid furrow ingression. Supporting this idea, the dominant negative N-terminal Bro1 domain that stabilizes the VLF causes delayed or failed cytokinesis (Figure 4).

While our *in vitro* studies support a role for the VLF in attachment and cytokinesis, the VLF could have another important role *in vivo*. The VLF is at the host-parasite interface and can fully encircle trophozoites (Figure 1B). It is known that *Giardia* secretes a number of effector proteins to both systemically modulate the immune response but also shorten microvilli directly under the trophozoites to potentially reduce the overall trophozoite profile and exposure to shear forces (52–55). The gasket-like VLF could be important for maintaining the concentration of specific secreted effector proteins beneath trophozoites. The microtubule sheets that form the ventral disc is too dense for vesicles to pass through, but at the center of the ventral disc there is a region lacking microtubules known as the bare area. Supporting the idea that there could be trafficking directed beneath the cell, there is a high concentration of vesicles associated with the bare area of the ventral disc (56, 57). Endocytosis markers indicate that internalization is occurring here (10, 57, 58), but whether directed secretion occurs at the bare area has not been studied. Newly developed genetic tools and reporter strains will allow for *in vivo* virulence studies to be performed in the future (59–61).

In summary, we identified Flangin, a protein that complexes with *Gl*Actin and the combination of these proteins form the backbone of the VLF. In parallel to the role of Rho GTPases regulating lamellipodia formation, *Gl*Rac regulates VLF formation. In the future, we will work to identify the molecular effectors that recruit *Gl*Actin and Flangin to the VLF, which is dynamically regulated over the cell cycle. Live imaging revealed that the VLF grows to ~1 μm in width after cell division then remains static in interphase, grows during mitosis, and is resorbed during cytokinesis. We experimentally confirmed that the VLF contributes to attachment and propose that this function is particular important during mitosis when the VLF is at its maximal width and the ventral disc is sequentially disassembled (11, 30). We also found that VLF has an innovative secondary role as a membrane reservoir to support *Giardia’s* rapid cytokinesis. Together, these observations point toward the biological importance of *Giardia* minimizing time detached for its parasitic lifecycle.

## Material and Methods

### Strain and culture conditions

*Giardia* strain WB Clone 6 (ATCC 50803) was cultured as in (62).

### Vector Construction

All constructs used in this study were made using standard techniques, see Table S2 for sequences and workflow. Construction of the morpholino sensitive HA-*Gl*Rac is described in reference (63).

### Immunoprecipitation

500 mL of *Giardia* wildtype and Flangin-3HA cell cultures were grown for 3 days, then iced for 2 hours to detach and pelleted at 1500xg at 4 °C. Cells were then washed twice in HBS with 2XHALT protease inhibitors plus 10 μM chymostatin, 1μM leupeptin, and 1 μM E64. Each pellet was resuspended to a final volume of 1.2 mL. Then 100 mM DSP (dithiobis(succinimidyl propionate)) in DMSO was added to a final concentration of 1 mM and incubated at room temperature for 30 minutes. The reaction was quenched for 15 minutes with 20 mM Tris pH 7.4. Cells were then pelleted by spinning for 7 minutes at 700xg and resuspended in 350 μL lysis buffer (80 mM KCl, 10 mM imidazole, 1 mM MgCl2, 1 mM EGTA, 5% Glycerol, 20 mM HEPES, 0.2 mM CaCl2, 10 mM ATP, 0.1% Triton X-100, 250 mM NaCl, pH 7.2). Cells were lysed by sonication and cleared with a 10 minute spin at 10,000xg. A volume of 17.5 μL of equilibrated EZview Red Anti-HA Affinity gel (Sigma) was added to each tube of lysate, then incubated at 4°C with end-over-end mixing for 1 hour. Beads were then spun at 8,200xg for 30 seconds and the supernatant was discarded, followed by a total of three washes with 750 μL of lysis buffer. Each wash consisted of end-over-end rotation for 5 minutes followed by a 30 second spin at 8,200xg. Protein was then incubated with 50 μL of 8 M Urea at RT for 20 minutes to elute bound proteins without releasing heavy or light chains from the anti-HA antibody. Sample buffer was then added to eluted samples, boiled for 5 minutes, then run on 12% SDS-PAGE gel followed by Western blotting.

### Western Blotting

Multi-plexed Western blots were performed by transferring proteins to Imobilon-FL membrane in Running Buffer (0.025M Tris, 0.192 M Glycine with 20% methanol, the blot was blocked in TBS (20mM Tris pH 7.6, 143 mM NaCl) plus 0.2% Tween-20 and 5% nonfat dry milk. Primary antibodies (anti-*Gl*Actin (10), anti-acetylated tubulin 6-11-B1 (Sigma), or anti-HA HA7 (Sigma)) were incubated overnight at 1:3000 at 4°C. After three washes (5, 10, 15 minutes) the blot was incubated for 60 minutes with secondary Goat Anti-Rabbit-HRP (Bio-Rad, 1:7000) and Alexa 647 Goat-Anti-Mouse antibodies (Molecular Probes, 1:2500). After three washes with TBS-T the blot was imaged with a Chemidoc MP (Bio-Rad). Values were normalized against the indicated loading controls after measuring band intensity with the ImageJ multi-measure function.

### Morpholino knockdown

Knockdown experiments were performed as described in (64) using the actin morpholino oligonucleotide 5’ GCAGGGTTGTCGTCTGTCA-TTTAC 3’, Flangin morpholino oligonucleotide 5’CGAGAGCGCAAGGATTCTGATGCAT 3’, *Gl*Rac morpholino oligonucleotide 5’ TATCCTCATTTCCTGTACTAGTCAT 3’, and the standard control morpholino oligonucleotide 5’ CCTCTTACCTCAGTTA CAATTTATA 3’, sourced from Gene Tools LLC.

### Immunofluorescence and expansion microscopy

*Gl*Actin localization at the VLF was not apparent in our prior localization studies, fixation in culture tubes does not preserve display of the VLF after settling in random orientations on cover glass. To observe the VLF, we allowed live cells in growth medium to attach to poly-L-lysine coated cover glass before fixation. Fixed imaging was modified from (63) using starve and release cells (3.5 hours) to increase mitotic index (62). For expansion microscopy fixation and staining proceeded as in (11). The expansion steps were performed as described in (28, 65).

### Fluorescent live imaging

Cells were chilled with ice for 20 minutes to detach from the culture tube and then placed into an Attofluor cell chamber (Molecular Probes) and incubated in a GasPak EZ anaerobic pouch (BD) or a Tri-gas incubator (Panasonic) set to 2.5% O_2_, 5% Co_2_ for 1-2 hours at 37° C. Cells were then washed four times with SB5050 (0.1% K2HP04, 0.06% KH2P04, 1% glucose, 0.2% NaCl, 0.2% cysteine-HCI monohydrate, 0.02% ascorbic acid, 0.0228% ferric ammonium citrate, 0.05% bovine bile and 5% bovine serum, pH 7.1). Cells were overlaid with a mixture of 0.7-1% ultra-low gelling agarose (Sigma A2576) melted in HBS (137mM NaCl, 5mM KCl, 0.91mM Na_2_HpO_4_-heptahydrate, 5. 55mM Glucose, 20mM HEPES, and pH7) and diluted into SB5050, left at room temperature for 10 minutes to solidify the agarose. Imaging was performed under 2.5% O_2_, 5% CO_2_, and 37° C (Oko Lab Boldline CO_2_/O_2_). For imaging of mitotic cells, the cells were treated with 0.25μM Albendazole ~4 hours before being imaged to increase the mitotic index. For Halo-tag labeling cells were allowed to re-attach for 1 hour under hypoxic conditions and then incubated with 0.5 uM of Janelia Fluor 549 (Promega) for 15 minutes. Cells were then washed four times with pre-warmed growth media and then allowed to incubate another 30 minutes in growth media to allow diffusion of unbound intracellular dye out of the cell before proceeding to image as above. Time-lapse imaging was performed on a DeltaVision Elite deconvolution microscope (GE, Issaquah, WA) equipped with DIC optics, using a 100×1.4 NA or 60× 1.42 NA objective, and a PCO Edge 5.4 sCMOS camera (PCO-TECH Inc).

### Fluorescence Recovery after Photobleaching

Attofluor cell chambers were seeded with Flangin-mNG parasites as outlined above. Imaging was conducted using a FRAP-enabled DeltaVision Spectris, 488 nm solid-state laser, 100% laser power, 50 mW laser, and 25 ms stationary pulse. The first image was acquired approximately 2 ms after the bleach event. The first minute, images were acquired every 10 second for 1 minute, then every 20 seconds for the following 2 minutes, every 50 seconds over the following 6 minutes, and finally, every 100 seconds for the remainder of the experiment. Normalized mNG fluorescence recovery was calculated by subtracting the background noise from the ROI intensity measurement; the background-subtracted intensity measurement was then divided by a fluorescent control ROI intensity measurement to normalize for photobleaching due to imaging.

### Scanning electron microscopy

Performed as described in (30). Briefly, 3% glutaraldehyde in 0.1M cacodylate buffer was used for trophozoite fixation, and 1% OsO_4_ in 0.1M cacodylate buffer for postfixation. Washing steps included 5% sucrose in cacodylate buffer, and PBS. Dehydration proceeded in an ethanol series prior to critical point drying and gold coating. Imaging was performed on a JEOL 6300 scanning electron microscope.

### Focused Ion Beam-Scanning Electron Microscope Tomography

Trophozoites were allowed to attach to laser marked slides (LASERMarking, Munich, Germany) in a silicon anaerobic chamber at 37°C. Slides were rinsed with PBS, pH 7.2 and immediately fixed with 2.5 % (v/v) glutaraldehyde (Science Services GmbH, Munich, Germany) in 75 mM cacodylate (Sigma-Aldrich), 75 mM NaCl, 2 mM MgCl 2 for 30 min, followed by 3 washing steps in cacodylate buffer. Cells were stained with DAPI, sealed with a coverslip and Fixogum (Marabu GmbH, Tamm, Germany) to prevent drying during LM investigation on Zeiss Axiophot fluorescence light microscope. The positions of cells of interest (ROIs) were marked on a template, with the same coordinates. For documentation, epifluorescence, phase contrast and DIC images were taken to retrieve the ROIs in FIB/SEM. All other steps, i.e. postfixation, ultrathin embedding, mounting, imaging in a Zeiss Auriga FIB/SEM workstation operating under SmartSEM (Carl Zeiss Microscopy, Oberkochen, Germany) and image analysis using Amira^®^ (Thermo Fisher Scientific, USA) were carried out as described in (66).

### Cell attachment assays

Cells were depleted for either Flangin, *Gl*Actin, or *Gl*Rac (see above for morpholino protocol), allowed to recover for ~24 hours in a total volume of 13 mL of media before performing the attachment assays.

#### Basic Assay

Culture tubes were inverted several times and then the detached cells were decanted and counted with a MoxiZ (Orflo) coulter counter. The media was replaced with 13 mL of ice-cold media and then placed on ice for 30 minutes to detach the remaining attached cells. These cells were then counted as above. Three independent replicates of each cell line and control were analyzed.

#### Flow Chamber Assay

Microfluidic devices were fabricated using a silicon master, which was created using deep reactive ion etching. The silicon master contained a raised channel that had a rectangular cross-sectional area, with a length of 1 cm, a width of 1 mm, and a height of 115 μm. A polydimethylsiloxane (PDMS) (Sylgard 184, Dow Corning) mold was made using a 10:1 base to curing agent ratio, which was mixed for 5 minutes, degassed for 20 minutes, poured over the master, and placed in an oven for 10 minutes at 110 °C to cure. The mold was peeled away from the silicon master and holes were punched for the inlet and outlet of the channel.

Afterwards, inlet and outlet ports were created using a custom aluminum mold with pegs to insert the silicone tubing (0.040 inch ID/ 0.085 inch OD, HelixMark). Degassed 10:1 PDMS was poured into the preheated aluminum mold and cured at 110 °C for 1 hour. The inlet and outlet ports were aligned with holes on the top face of the microfluidic device and stuck together using uncured PDMS and 5 minutes of oven heat. Finally, the bottom face of the microfluidic device and a clean glass slide were plasma treated for 10 seconds and aligned together to create a water-tight seal between the two layers.

Trophozoites were chilled for 20 minutes on ice and then loaded into microfluidic devices placed in a GasPak EZ anaerobic pouch (BD Scientific) at 37 °C for 1 hour to allow re-attachment. Warm growth media was drawn into a disposable syringe (BD Scientific) and then loaded onto a syringe pump (Harvard Apparatus). For experiments, the flow rate was set to 20 μl/s for 30 seconds to clear swimming trophozoites and then ramped to 100 μl/s over 5 seconds and challenged for 60 seconds. The challenge assays were recorded at 2 FPS using a Nikon Eclipse Ti microscope with a 20× phase objective, Clara DR-1357 CCD camera (Andor), and a live-cell incubation chamber (In Vivo Scientific, Inc) to maintain a temperature of 37 °C. Trophozoite displacement was quantified starting at 35 seconds using the TrakMate ImageJ plugin via manual tracking over the first 10 seconds of 100 μl/s flow (67, 68).

### Statistical analysis

Statistically significant differences between KD and control groups were obtained using GraphPad Prism 7.02 software and unpaired t test with Welch’s correction for parametric data and the Mann-Whitney U test for non-parametric data.

## Supplemental Files

**Movie 1** Flangin dynamics during mitosis and cytokinesis. Note that in a portion of cells a cytoplasmic bridge can persist between daughter cells for tens of minutes (30).

**Movie 2** DIC imaging of cytokinesis and VLF regrowth. Video also shows the VLF remains size uniform after re-growth (filmed for >50 minutes).

**Movie 3** DIC imaging of morpholino control cell through mitosis and cytokinesis.

**Movie 4** DIC imaging of a Flangin depleted cell that restores VLF width before cytokinesis.

**Movie 5** A dominant negative Flangin_1-362_-mNG cell that fails to retract the VLF and fails cytokinesis. Note that this cell remains attached to the cover glass and the VLF width increases as filopodia like projections are formed.

**Movie 6** Dominant negative Flangin_1-362_-mNG imaged with DIC only to avoid oxidative stress. This completes cytokinesis at 8 minutes, the delay is associated with partial failure to resorb the VLF.

**Movie 7** Representative attachment assay under flow. Morpholino control on the left and Flangin knockdown on the right. Colored lines indicate the trajectory of individual cells beginning at 35 seconds when the flow rate peaks at 100 μl/s.

**Supplementary Table 1** Spreadsheet containing putative WH2 domain containing proteins.

**Supplementary Table 2** Spreadsheet containing primer sequences and construct design.

## Acknowledgements

We thank Renyu Li and Andrew Shelton for technical assistance, Prof. Gerhard Wanner LMU, Munich Germany for sharing his outstanding expertise on FIB/SEM tomography. This work was supported by NIH Grant 5R01AI110708 (A.R.P.), NSF Grant GRFP DGE-1256082 (W.R.H.), NIH Grant 1F32AI145111 (K.L.H.), Charles University grant PRIMUS/20/MED/008 (P.T.), Czech Republic MSMT-10251/2019 grant LTAB19004 (P.T), Burroughs-Wellcome Career Award at the Scientific Interface (J.C.V.), and NIH grant R01MH115767 (J.C.V).

---

**Figure S1.**
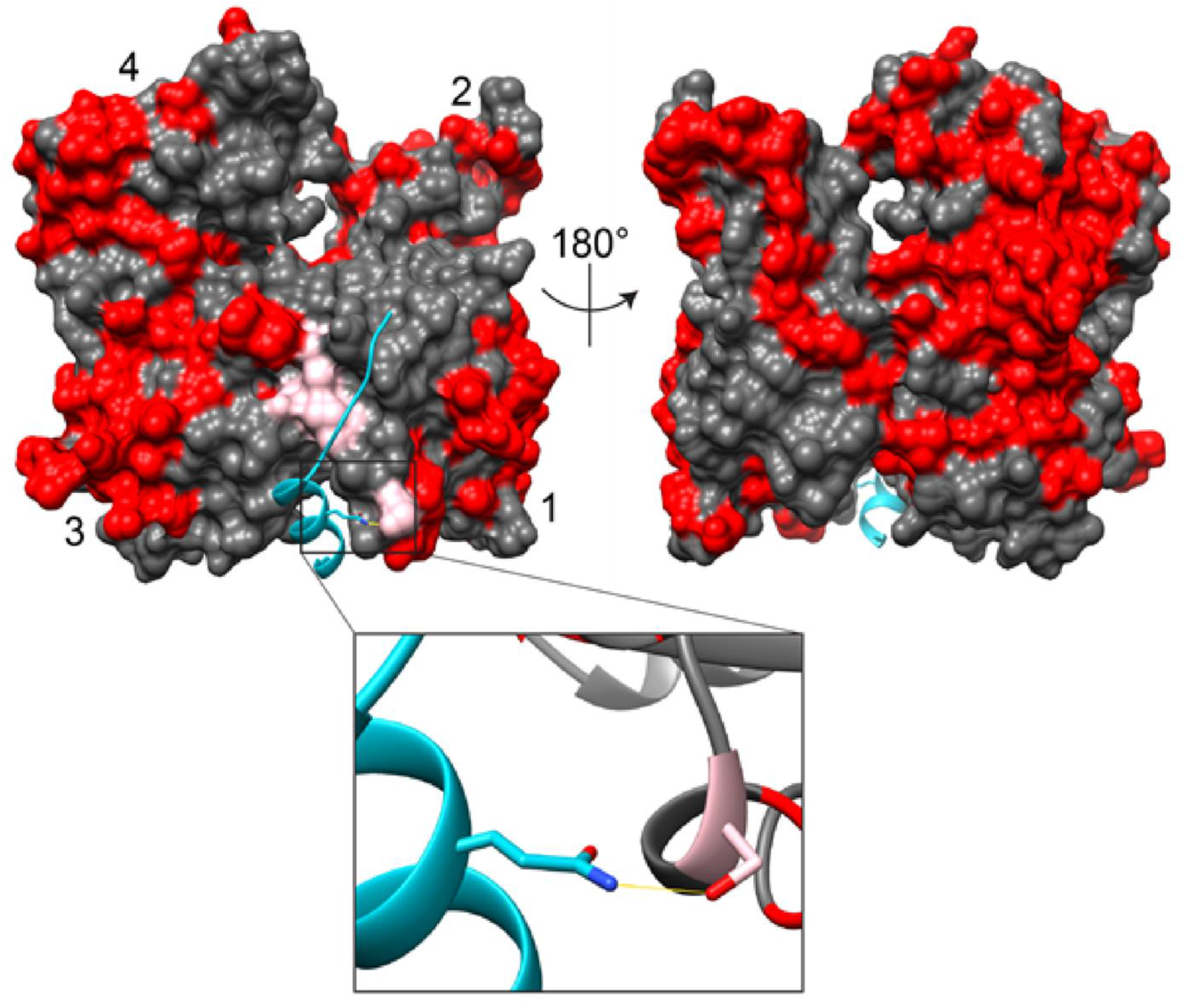
Variations in the amino acid sequence of *Gl*Actin are minimal within the WH2 binding region. Modeled surface of *Gl*Actin based on mammalian skeletal actin (PDB 2A3Z). Grey surface shows conserved residues, with red sections highlighting variations between skeletal actin and *Gl*Actin. Pink resides are those residues that differ and are within 5 Å of the bound WH2 domain of WASP (blue ribbon). *Inset*: Only one reside change, Ser350Ala, would cause the loss of a hydrogen bond between the WH2 domain and *Gl*Actin (ribbons with WH2 residue Gln437 and Ser350 shown as sticks). Residue 437 is not part of the conserved WH2 motif (16).

**Figure S2.**
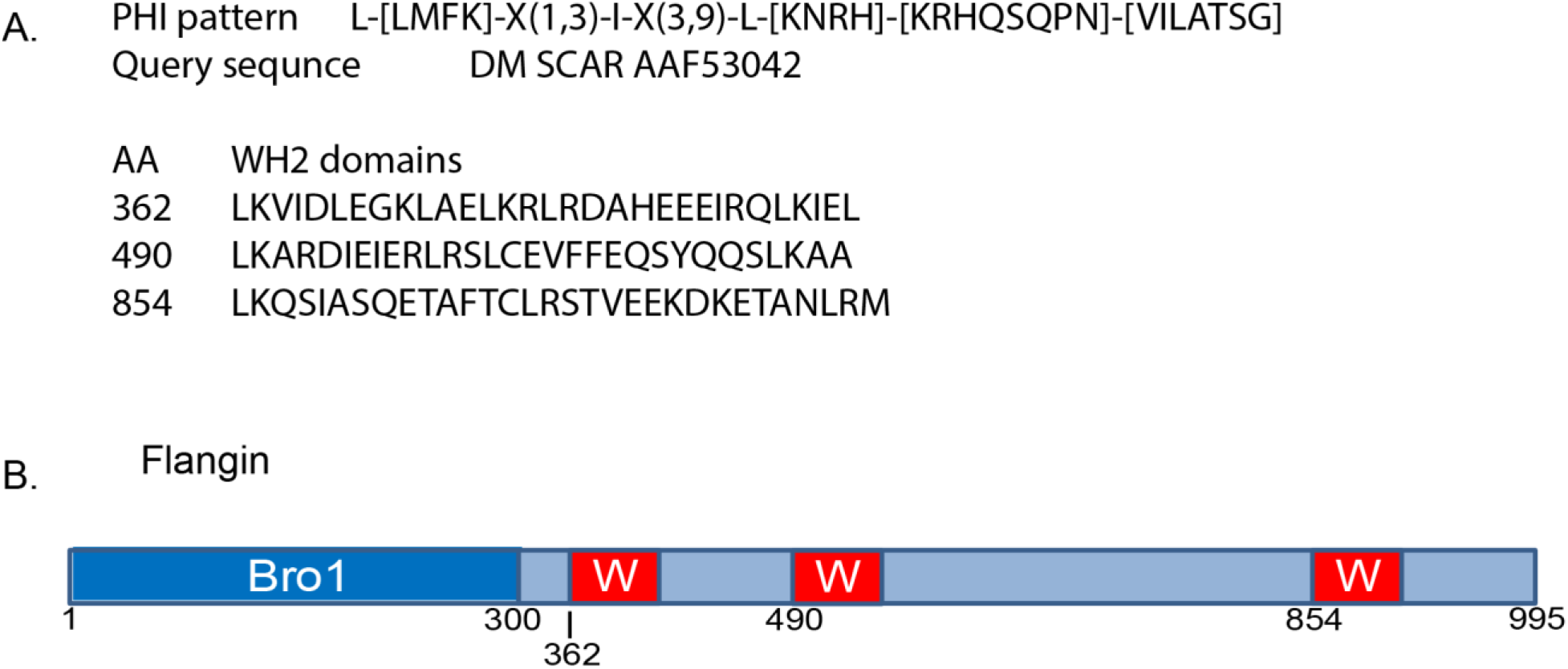
Diagram of Flangin domain organization. (A) PHI BLAST with DM SCAR AAF53042 as the query sequence identified three WH2-like domains in Flangin (GL50803_7031). (B) Diagram of Flangin domain organization with amino acid positions indicated.

**Figure S3.**
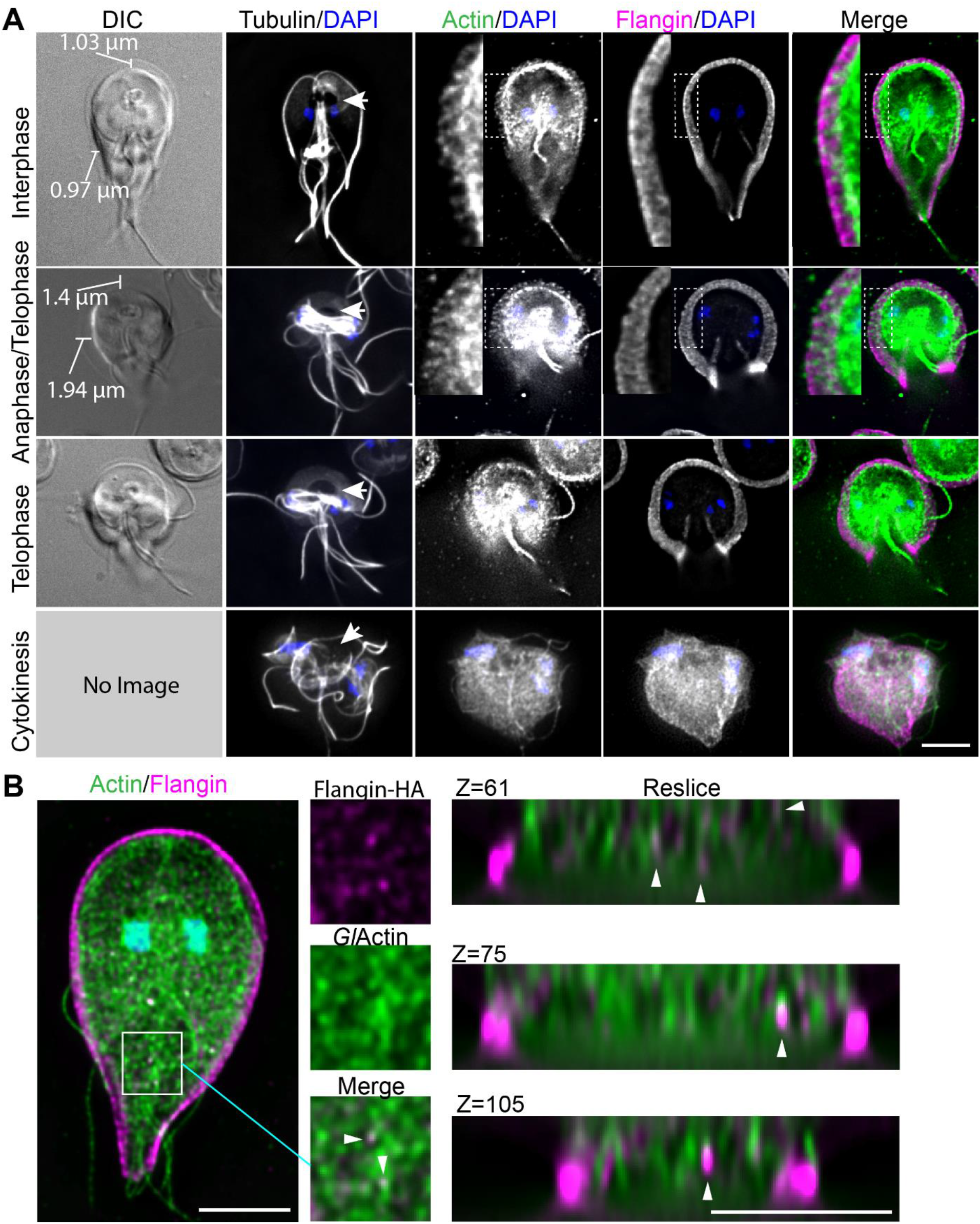
The VLF grows in width before ventral disc disassembly in telophase. Immunofluorescence localization of tubulin, *Gl*Actin (green), Flangin-HA (magenta), and DNA (blue), throughout the cell cycle. *Gl*Actin and Flangin localized to the VLF during interphase and mitosis. The VLF was wider in mitosis corresponding to when the ventral disc begins conformational changes and disassembly as indicated by the size of the bare area (Arrows). This microtubule free region grows as the ventral disc is disassembled in telophase. Ultimately the ventral disc opens up and completely disassembles to permit furrow progression during cytokinesis. The insets show magnified views of *Gl*Actin and Flangin in the VLF. The VLF is resorbed during cytokinesis, which corresponds to when the cells begin to swim apart to generate membrane tension. Note that the interphase and mitotic cells are partial projections optimized to show *Gl*Actin localization in the VLF, while the entire Z-stack was projected for the cells in cytokinesis to show that the VLF has been resorbed. Also see Figure 1 and Movie 1 of (11), which shows microtubules and VLF in live cells. (B) Flangin (magenta) can be seen at the ends of *Gl*Actin filaments (green) in maximal projections and re-sliced image stacks. Scale bar= 5 μm.

**Figure S4.**
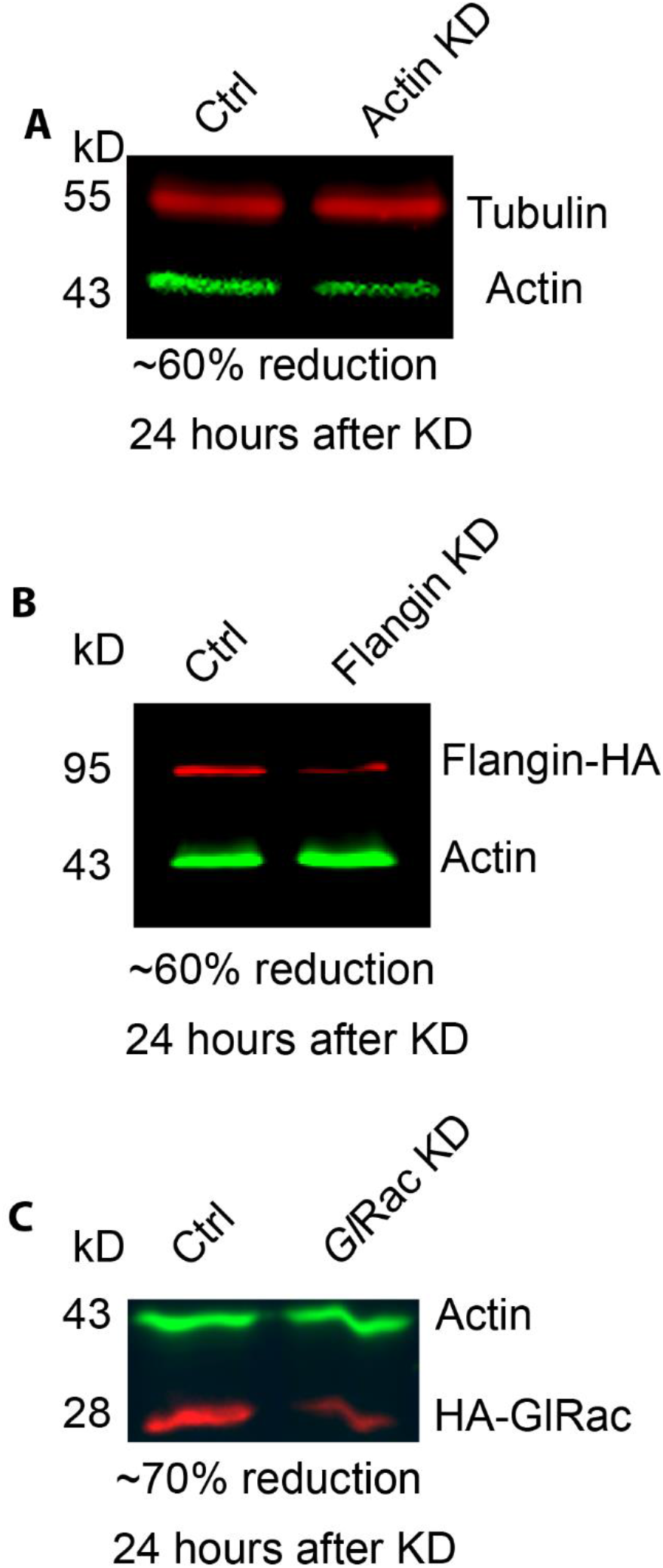
Morpholino efficacy. (A) Standard control morpholino versus translation blocking anti-*Gl*Actin treatment specifically reduced *Gl*Actin by ~60% as observed by Western blotting. Protein levels were normalized using tubulin as a loading control. (B) Standard control morpholino versus translation blocking anti-Flangin morpholino reduced Flangin levels by ~60% based on Western Blotting. Protein levels were normalized using actin as a loading control (C) Standard control morpholino versus translation blocking anti-*Gl*Rac morpholino reduced *Gl*Rac levels by ~70%. Protein levels were normalized using actin or tubulin as a loading control. All blots were performed 24 h after morpholino treatment, matching the timing of live and fixed cell experiments.

**Figure S5.**
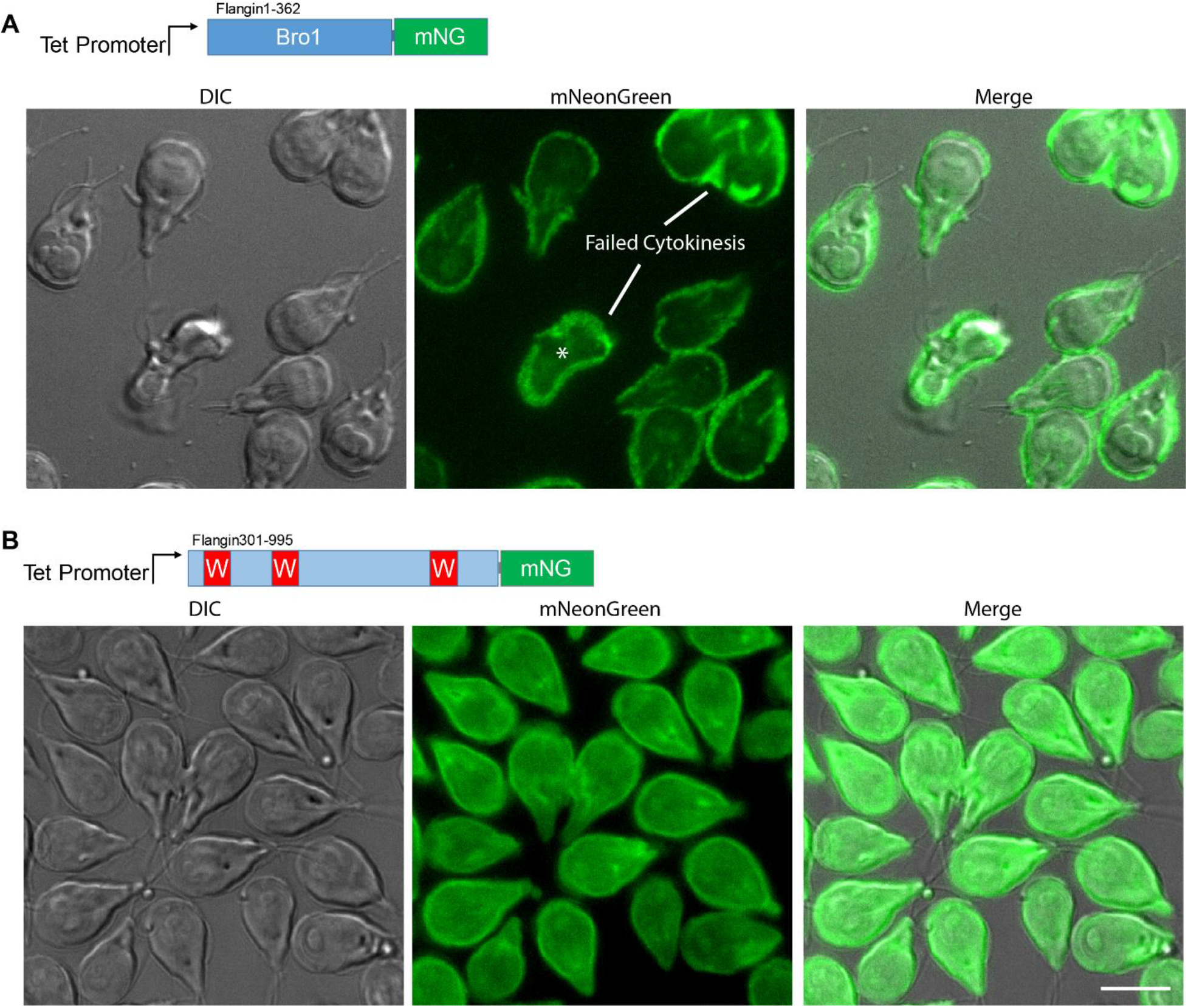
Localization of the N-terminal Bro1 domain and C-terminal WH2 domain fragments in live cells. (A) The N-terminal Bro1 fragment is recruited to the same regions as full length Flangin. Expression of this fragment results in failed cytokinesis, asterisk marks cell actively trying to divide that failed to break down the VLF. (B) The C-terminal region containing putative WH2-like domains is not specifically recruited to the VLF. Scale bar= 10 μm.

## Notes

### Competing Interest Statement

The authors have declared no competing interest.

